# OTTR-CLASH: improved biochemical and bioinformatic identification of Argonaute 2-mediated microRNA-target RNA interactions

**DOI:** 10.64898/2026.07.14.738487

**Authors:** Paul D. Kaufman, Haibo Liu, Kai Hu, Lucas Ferguson, Kathleen Collins, Lihua Julie Zhu, Thoru Pederson

## Abstract

Various methods have detected miRNA-target interactions via immunoprecipitation of UV-crosslinked Argonaute ribonucleoprotein complexes, followed by intermolecular ligation of bound miRNAs to target strands, forming chimeric RNAs. To date, these methods have relied on conventional viral reverse transcriptases (RTs) to generate cDNAs for sequencing. However, crosslinked RNAs often retain adducts after purification, which can make them poor templates for viral RTs. Here, we adapted OTTR (Ordered Two-Template Relay) techniques to generate cDNAs from Ago2-bound RNAs. OTTR makes use of a modified retroelement-encoded RT, which is strongly processive even on templates with modifications or adducts. We show that this “OTTR-CLASH” method increases the frequency of generating chimeric RNAs compared to previous methods. We also developed an improved bioinformatic pipeline for analysis of these data, and we use this to catalog miRNA-target interactions not previously described in the literature.

**Graphical Abstract:** 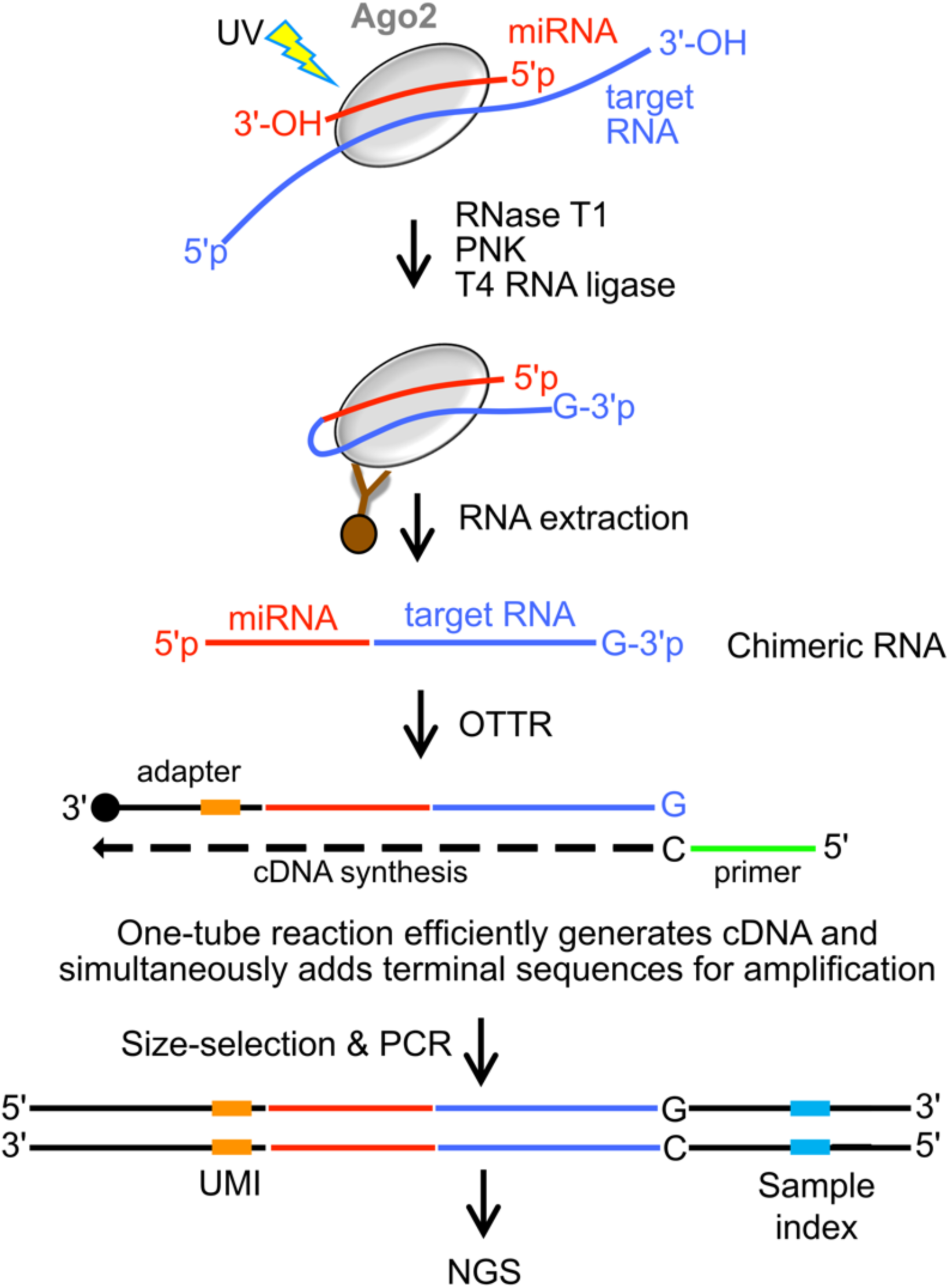

## INTRODUCTION

MicroRNAs (miRNAs) are small noncoding RNAs with lengths between 17 and 25 nucleotides (nt) (1, 2). They are key regulators of many cellular processes and are critical for normal development, physiology and health in most eukaryotic organisms (3, 4). Aberrations in the biogenesis, expression level and activity of miRNAs are frequently reported in many human diseases (5, 6), further supporting their contribution to a wide range of biological processes.

miRNAs regulate cellular fate and/or homeostasis (7–9) through binding to target RNAs such as messenger RNAs (mRNAs), long noncoding RNAs (lncRNAs), and pseudogene transcripts (10–12). These binding interactions can regulate target RNA stability and/or translational rate (3, 13, 14) . miRNAs execute their functions after being loaded into the ‘RNA-induced silencing complex’ (RISC). RISC is a multisubunit enzyme that performs the final miRNA maturation step and is essential for recognition of specific target RNAs (15, 16). The key RISC subunits are Argonaute (Ago) proteins, of which there are four in humans (17) . Ago2 is the only human Argonaute protein with endonucleolytic RNA cleavage activity (“slicer”) for maturation of miRNAs (18, 19). Ago2 is also the only Argonaute essential for early mouse development (20). Therefore, experimental detection of functional miRNA-target interactions often begins with isolation of Ago2-containing complexes bound to RNAs (21, 22). Current bioinformatic calculation methods alone cannot yet completely predict all physiologically relevant miRNA-target interactions, despite inclusion of conservation and sequence context information (23–26). This uncertainty is exacerbated by the ability of Ago proteins to engage targets via “non-canonical” basepairing interactions (27–29). Therefore, an experimental approach to identifying transcriptome-wide miRNA-target interactions is much needed.

Previous studies have described immunoprecipitation (IP) of UV-crosslinked Ago2 complexes, followed by intra-complex ligation of miRNAs to their targets and reverse transcription of the resulting hybrid RNAs prior to library preparation for deep sequencing (30–36). Critical limitations common to all previous methods include the inherent low efficiency and high sequence bias of adapter ligation reactions required for Illumina-compatible library preparation (37–40). Furthermore, the use of UV crosslinking results in RNA molecules that contain bulky adducts (41) that block polymerization during reverse transcription by conventional retroviral RTs (42).

In this study, we introduced an improved methodology for generating Ago-bound chimeric RNAs. We replaced both the conventional RT step and the ligation-dependent NGS library preparation with a single-tube reaction called OTTR (“Ordered Two-Template Relay”). Recent work established OTTR as a streamlined and efficient method for the generation of cDNA libraries for high-throughput sequencing (43, 44), particularly when the input RNA is of low abundance and/or chemically modified (45, 46). The improved conversion of RNAs to cDNAs in OTTR results from the enzymatic properties of a silkworm non-LTR retroelement reverse transcriptase called BoMoC (47). BoMoC is more processive on modified RNA templates than conventional reverse transcriptases. Furthermore, during the cDNA synthesis reaction it also attaches adapter sequences, eliminating the need for inefficient ligation steps (43, 44). We reasoned that this enzyme would be well-suited to analysis of miRNA-target hybrids, because the hybrids are generated from UV-crosslinked samples (Fig. 1A).

**Figure 1:**
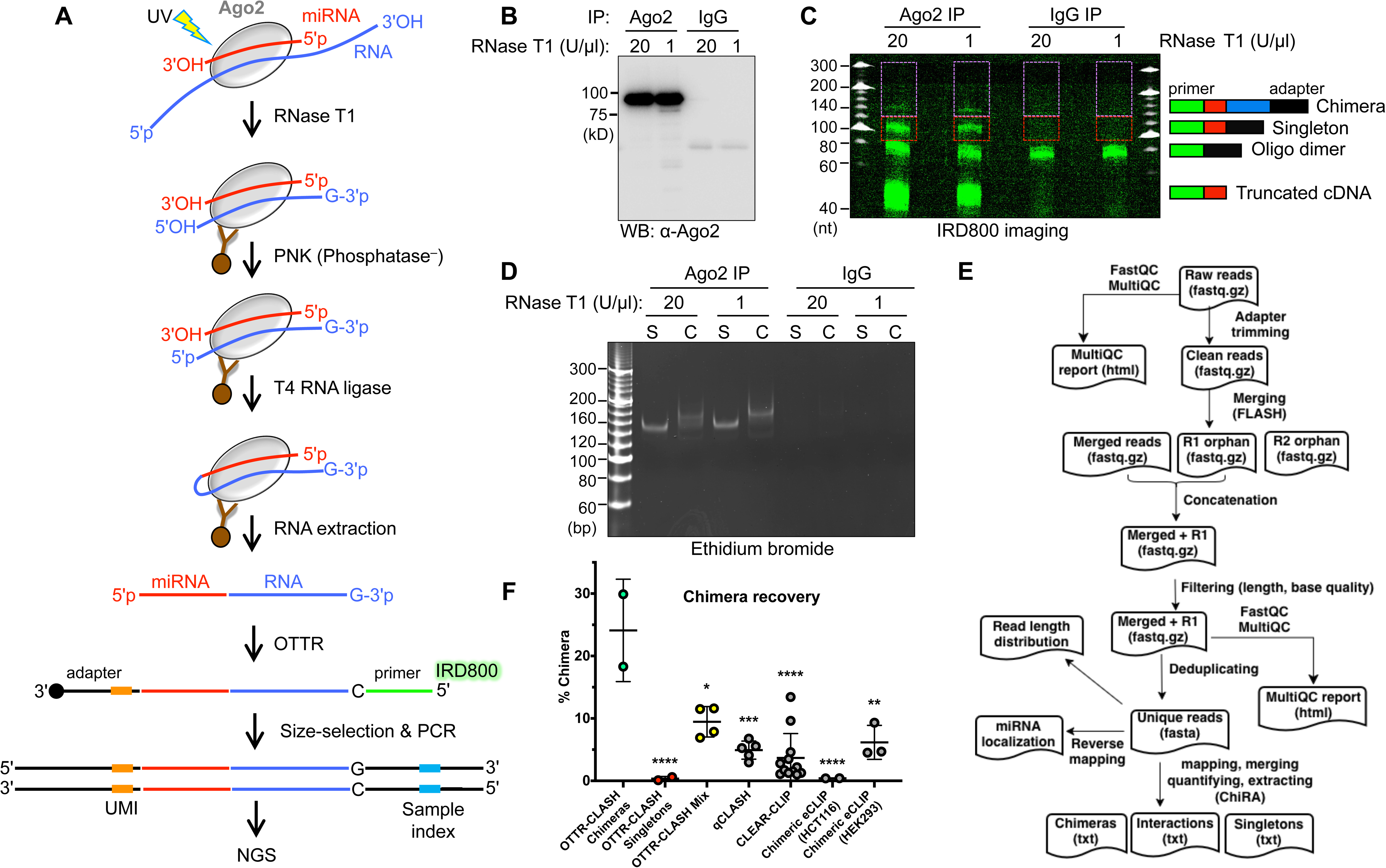
OTTR-CLASH detects Ago2-bound chimeric RNAs at a higher frequency than previous methods. A. Steps in the OTTR-CLASH protocol. “primer” indicates the IR800-labeled primer duplex, colored in green. The miRNA (red) and target RNA (blue) strands bound to Ago2 become a chimeric molecule after ligation. The adapter (black) is illustrated with a 3’ chain terminating group (round circle), which prevents formation of higher order products during the OTTR reaction. B. Western blot confirmation of Ago2 immunoprecipitation from HeLa-S3 whole cell extracts using monoclonal antibody 4F9. IgG was used as the negative control antibody. Proteins were detected using anti-Ago antibody 11A9. C. Chimeric cDNAs were generated by OTTR-CLASH, separated via denaturing 8% PAGE and detected by IR800 imaging. Product structures are illustrated on the right, with the color coding the same as in panel A. The “Chimera” (>∼140nt) and “Singleton” bands that were cut from the gel are shown by magenta and red rectangles, respectively. Note the greater amount of material in the chimera bands in the Ago2 IP samples compared to the IgG controls. “Oligo dimer” denotes products formed upon joining the primer duplex and adapter without insert, and “Truncated cDNA” denotes reaction intermediates formed without joining to the adapter oligo. D. Native PAGE analysis of AMPure-purified sequencing libraries. “S” Singleton-enriched material amplified from the lower bands in panel C. “C” indicates chimera-enriched material enriched from upper bands. Note that Ago2 IP samples (lanes 1-4) produced visible products while IgG samples (lanes 5-8) did not. Two concentrations of RNase T1 (20 and 1 U/µl) were used to prepare the whole cell extracts. E. Bioinformatics workflow for the analysis of chimeric reads. F. Comparison of chimeric read percentages in OTTR-CLASH experiments with those in previously described methods. OTTR-CLASH samples generated by gel separation into “Chimeras” (green dots) and “Singletons” (red dots) were considered separately from samples in which a single gel slice containing all products with inserts was isolated (“Mix”, yellow dots). p-values from one-way ANOVA comparisons between the OTTR-CLASH Chimeras and each other data set are presented as asterisks (*, < 0.05; ** < 0.01; *** < 0.001, **** < 0.0001).

We fused OTTR with established steps for amplifying miRNA-target hybrids (*e.g.* as in methods known as qCLASH (quick Crosslinking, Ligation And Sequencing of Hybrids (33, 34)), and Chimeric-cCLIP (36). We now present this ‘OTTR-CLASH’ workflow, which increases the recovery of miRNA-target hybrids compared to previous methods. Furthermore, we developed an improved bioinformatic analysis package for the mapping of the chimeric reads generated in these experiments. Our biochemical and bioinformatic pipeline detected a wide variety of interactions not previously documented in the literature. The ease of use of this protocol should make it generally useful for all investigators of miRNA-target interactions.

## Materials and Methods

### Cell culture

HeLa-S3 cells (ATCC CCL-2.2) were cultured and maintained in suspension in spinner flasks (Bellco, 3622129) containing Leibovitz’s L-15 medium (ATCC 30-2008) supplemented with 5% horse serum (Gibco, 16050122) and 1% Pen/Strep in atmospheric air at 37°C with constant stirring. Cells were maintained at 3 – 5 × 10^5^ cells/ml (48, 49). The cultures were provided new media every 48 hours.

For storage, HeLa-S3 cells were frozen in 90% fetal bovine serum (FBS) + 10% DMSO (Sigma, 34869-100ML). For recovery of frozen cells, they were thawed in Minimum Essential Medium (Thermo, 11095072) containing 5% FBS and 1% Pen/Strep, collected via centrifugation (500 × g for 5 min at 4°C), and plated onto 10 cm plates. After this initial culturing, cells were trypsinized and transferred to a small spinner flask (Bellco – 500 ml) in at least 200 ml complete Leibovitz’s L-15 medium at an initial density of 1 – 1.5 × 10^5^ cells/ml and maintained up to 5 × 10^5^ cells/ml. For large-scale experiments, 3 – 4 liters of this final suspension of growing cells were used. Samples were diluted 1:1 in 0.4% Trypan Blue (Sigma, T8154) and counted on a hemocytometer to check viability and count the cells in suspension (50).

### UV-crosslinking for OTTR-CLASH

HeLa-S3 cells in suspension (up to 4 L) were split into several 750 ml centrifuge bottles (Beckman, 356855), collected by centrifugation at 500 × g for 5 min, and supernatants were discarded. Pelleted cells were washed once with ice-cold PBS (∼ 8× packed cell volume) to remove any residual media, to avoid trace amounts of phenol red interfering with UV irradiation (51). Washed cells were resuspended in ice-cold PBS at a density not greater than 50 million cells per ml cold PBS for crosslinking. This cell suspension was distributed into several 15-cm non-tissue culture-treated plastic Petri dishes (20 ml cell suspension per plate), placed on ice (with the lid open), and irradiated with 254 nm light with a UVP CL-1000 UV crosslinker. To maximize cell exposure to the UV irradiation, plates were crosslinked once at 400 mJ/cm^2^, rotated 180° and swirled, and crosslinked again at 200 mJ/cm^2^ for a total dose of 600 mJ/cm^2^ (34). The UV-irradiated cells were transferred from 15-cm plates into 50-ml conical tubes on ice, then pelleted by centrifugation at 200 × g for 5 min. In cases where there was significant adhesion of cells to the plastic after crosslinking, after the initial cell transfer 10 ml cold PBS was added to the attached cells, which were then dislodged with plastic cell lifters (Cell-treat, 229306) and added to the collected cell suspension. The UV-crosslinked cells were collected by centrifugation at 500 × g at 4°C for 5 minutes and the supernatant was removed via vacuum aspiration. Pelleted cells were then stored at -80°C until the time of next experimental step. 20 or 50 million cells per sample were used for each total RNA extraction or OTTR-CLASH experiment, respectively.

### Cell lysis

To make cell extracts for OTTR-CLASH experiments, ‘complete RIPA buffer’ was prepared by adding 11 µl Protease Inhibitor Cocktail III (Sigma, 539134-1ML), and 5 µl 0.1 M DTT (Thermo, R0861) per 1 ml RIPA (Thermo, 89901; 25 mM Tris-HCl pH 7.6, 150 mM NaCl, 1% NP-40, 1% sodium deoxycholate, 0.1% SDS). After mixing, the buffer was kept on ice prior to cell lysis.

For each lysate, 100 million frozen, UV-irradiated cells were lysed by addition of 5 ml pre-chilled complete RIPA buffer and mixing by pipetting on ice, followed by incubation on ice for 20 minutes in multiple 2-ml LoBind Eppendorf tubes (Eppendorf, 022431048). Lysates were then treated with Turbo DNase (10 µl per 1 ml lysate, (2 U/µl) (Thermo, AM2239)) and Murine RNase inhibitor (22 µl per ml lysate, (NEB, M0314S)) for 5 minutes at 37°C with shaking at 1000 rpm in an Eppendorf thermomixer. Digested lysates were cleared by centrifugation at 21,000 g for 5 minutes at 4°C and supernatants placed in fresh tubes and immediately treated with RNase T1.

### RNase T1 digestion

Digesting full-length Ago-associated transcripts to footprint-sized fragments improves the precision of mapping interactions between miRNAs and their targets (32). To do this, we used RNase T1, which was previously used in other methods to characterize Ago-associated RNAs (52–54). RNase T1 specifically digests single-stranded RNAs on the 3’ side of guanidine (G) residues, making it ideal for subsequent OTTR reactions that use a primer duplex with a single cytosine nucleotide overhang to initiate polymerization (43, 44). DNase-treated, cleared cell lysates were treated with RNase T1 (1000 U/µl, Thermo, EN0542) for 15 min at room temperature. To stop RNase T1 digestion, samples were placed on ice (for ≥ 5 minutes). Lysates were then divided into two samples, to perform both experimental (anti-Ago2) and control (IgG) immunoprecipitations.

Like other nucleases, the activity of RNase T1 may change in different types of biological samples (55, 56). Therefore, we tested multiple RNase T1 concentrations. ≥ 20 µl samples of the RNase T1-treated extracts were reserved as input samples for Western confirmation of the immunoprecipitation (IP) step (Fig. 1B).

### Ago2 Immunoprecipitation

For each sample, we used a total of 4 mg total protein lysate, together with 300 µl bead slurry of Dynabeads M-280 Sheep Anti-Mouse IgG (10 mg/ml, Invitrogen 11202D, hereafter called ‘M-280 beads’). We chose M-280 beads rather than Protein A or Protein G beads because M-280 beads do not require use of a bridge antibody for efficient IP of Ago proteins (34).

For each IP, three separate 100 µl aliquots of the Dynabead slurry were each transferred to their own 2-ml LoBind Eppendorf tube. Tubes were placed on a magnetic stand (DynaMag-2, Invitrogen, 123.21D) to separate the beads from the solution. For all washing steps, beads were first resuspended in solution of the magnetic stand, then placed onto the stand for removal of the supernatants via pipette-aspiration. Beads were washed twice with ice-cold, complete bead coating buffer (500 µl 50 mM Tris-HCl (pH 7.4), 100 mM NaCl, 1% NP-40, 0.1% SDS and 0.5% sodium deoxycholate supplemented with 5.5 µl Protease Inhibitor Cocktail III and 11 µl Murine RNase inhibitor). After washing twice, beads in each tube were resuspended in 100 µl complete bead coating buffer and 2 µg mouse anti-human Ago2 monoclonal antibody 4F9 (Santa Cruz, sc-53521) or normal mouse IgG as a negative control (Santa Cruz, sc-2025) per 100 µl initial bead volume were added. After pipetting to mix, samples were incubated for 45 minutes at room temperature with rotation. Antibody-coated beads were magnetically separated and washed twice with 500 µl ice-cold bead coating buffer (without protease and RNase inhibitors).

We observed that IPs worked best if the extract volume did not exceed 500 µl per 100 µl conjugated beads. Use of less than 1 µg antibody per ml lysate would also result in low IP efficiency.

500 µl cleared RNase T1-treated lysate (∼ 1.33 mg protein) was added to the antibody-conjugated magnetic beads in each 2-ml LoBind Eppendorf tube. Thereafter, beads were mixed by pipetting, incubated for 4 hours to overnight at 4°C with constant rotating, and then collected on the DynaMag-2 rack. The flow through (FT) supernatants were removed, with a ≥ 20 µl sample of the FT saved for Western blot-confirmation of the IP.

### Modifying terminal ends of the Ago-bound RNAs prior to ligation on beads

#### A) Stringent washing of the Ago2 complexes

After removing flow-throughs from the beads, beads were washed three times with 500 µl ice-cold High Salt Wash buffer (50 mM Tris-HCl (pH 7.4), 1 M NaCl, 1 mM EDTA, 1% NP-40, 0.1% SDS and 0.5% sodium deoxycholate) to reduce non-specific binding of lysate materials to the antibodies and/or the beads (57). Beads were then washed three times with 500 µl ice-cold Low Salt Wash buffer (20 mM Tris-HCl (pH 7.4), 10 mM MgCl_2_, 0.2% Tween-20 (Promega, H5152) and 5 mM NaCl).

#### B) 5’-end phosphorylation of the Ago2-bound RNAs

RNase T1 digestion leaves RNAs with a 5’-hydoxyl (5’-OH) group and a 2’ – 3’-cyclic and/or 3’-phosphate (3’P) group (34, 58). To favor ligation of miRNA 3’ ends to the 5’ termini of target RNAs, 5’-phosphorylation of the target RNAs was performed with a phosphatase-minus version of polynucleotide kinase (PNK), thereby minimizing dephosphorylation of target RNA 3’ ends (Figure 1A) [78, 83]. Beads were resuspended in 100 µl PNK reaction solution (1x PNK buffer (NEB), 1 mM ATP (Thermo, R0441), 20 mM NaCl (Invitrogen, AM9760G), 0.4 U/µl murine RNase inhibitor and 0.3 U/µl T4 PNK (phosphatase minus, NEB M0236S)). Samples were briefly mixed by pipetting to resuspend the beads, then incubated at 37 °C for 20 minutes with constant shaking at 1200 rpm.

### Ligation of miRNAs to their targets

After the PNK reaction, supernatants were magnetically separated from the beads and discarded. Beads were then washed once with ice-cold High Salt Buffer (500 µl) and three times with 500 µl ice-cold Low Salt Wash Buffer. While the beads were resuspended in their last washing solution on ice, ligation reaction components were assembled on ice (1x RNA ligase buffer (NEB, M0437M), 0.02% Tween 20 (Promega, H5152), 3% DMSO (Sigma, D8418-50ML), 1 mM ATP (Thermo, R0441), 15% PEG8000 (NEB), 0.67 U/µl Murine RNase inhibitor (NEB, M0314S)). 270 µl of the ligation mix was added per sample, and then T4 RNA Ligase enzyme (NEB, M0437M, high concentration) was added separately to each sample to a final concentration of 2.4 U/µl. Reactions were then incubated at room temperature for 1 – 2 h with rotation.

### 3’-end dephosphorylation of miRNA-target hybrid RNAs on beads

After ligation, supernatants were magnetically separated from the beads and discarded. Beads were then washed once with 500 µl ice-cold Low Salt Wash Buffer, once with 500 µl ice-cold High Salt Buffer, and twice with 500 µl ice-cold Low Salt Wash Buffer. To make the ligated miRNA-target hybrid RNAs suitable templates for the BoMoC enzyme, their 3’-end phosphate groups resulting from RNase T1 digestion must be converted into hydroxyl 3’-OH groups (44). Therefore, dephosphorylation using a combination of alkaline phosphatase (FastAP, Thermo, EF0651) and PNK (NEB M0201L, phosphatase-plus) was performed. Beads were resuspended in 50 ul FastAP reaction mix (1x Fast AP buffer (Thermo, EF0651), 1.6 U/µl Murine RNase inhibitor, 0.08 U/µl Turbo DNase (Thermo, AM2239), 0.06 U/µl Fast AP enzyme) and incubated at 37°C for 10 min with 1200 rpm shaking. Then, 150 µl of PNK mix (1x PNK buffer, 0.20 U/µl T4 PNK, phosphatase-plus (NEB M0201L) was added to the 50 µl FastAP reactions. Samples were incubated at 37°C for 20 min with 1200 rpm shaking in an Eppendorf Thermomixer.

### RNA extraction from beads

#### A) Bead washing

After the FastAP and PNK reactions, supernatants were separated magnetically and discarded. Beads were washed once with 500 µl cold High Salt Wash buffer, and then three times with 500 µl Low Salt Wash buffer.

#### B) Western blot confirmation of IP

In the last washing step, when the beads are resuspended in 500 µl Low Salt wash buffer, 10% of the beads in solution (*i.e.,* 50 µl out of 500 µl) was transferred into new 1.5 ml tubes for Western blot analysis to check the IP efficiency. The remaining amount (450 µl out of 500 µl) was subjected to proteinase K digestion followed by RNA extraction and cleanup (Steps C – D, below).

For preparing the eluates for Western blot analysis, the 50-µl bead solution was first placed on the magnetic rack and the supernatant was discarded. Then, 30 µl of an elution mixture containing 20 µl Low Salt Wash buffer, 3 µl 1M DTT and 7.5 µl NuPAGE LDS Sample Buffer (4×) (Invitrogen, NP0007) was added per sample. Samples were warmed at 70°C for 10 minutes with 1200 rpm shaking to facilitate dissociation of proteins from the beads. After 1 minute cooling samples on ice, samples were placed on the magnetic rack and their supernatants were saved and stored at -20°C until SDS-PAGE analysis. 15 µl of the 30 µl eluted samples were used for Western analysis, using rat anti-human Ago2 antibody, clone 11A9 (Sigma, MABE253), HRP-conjugated Goat Anti-Rat IgG, (Jackson Immuno-Research Laboratories, 112-035-008) and ECL detection (Thermo).

#### C) Proteinase K digestion

The remaining 90% of the Ago-antibody-bead complexes (450 µl total volume) was used for RNA extraction. Beads were placed on the DynaMag-2 magnetic rack and the supernatant was discarded. Beads were placed on ice and a mixture containing 97.5 µl 1x proteinase K buffer (100 mM Tris-HCl (pH 7.4), 50 mM NaCl, 10 mM EDTA and 0.2% SDS) and 22.5 µl proteinase K enzyme (NEB, P8107S, 800U/ml) was added to each sample. Samples were first incubated at 37°C for 20 minutes, and then 20 minutes at 50°C.

#### D) RNA cleanup with Phenol/Chloroform/Isoamyl alcohol (PCI)

The crude RNA samples were further purified and concentrated via organic extraction and alcohol precipitation. First, the 120 µl proteinase K-digested sample was adjusted to 400 µl by adding nuclease-free water (280 µl), then 500 µl PCI (VWR V0966, pH = 4.5) was added and samples were vigorously shaken (1400 rpm) at room temperature for 8 minutes. Samples were centrifuged at 4°C for 10 minutes at 18,000 g, and the upper aqueous phase was transferred into a new 1.5 nuclease-free Eppendorf tube on ice. To each separated aqueous phase, a mixture containing 1000 µl isopropanol-ethanol (1:1), 40 µl 3M sodium acetate (pH = 5.2) (Sigma, 567422-100ML) and 4 µl glycogen (20 mg/ml (Thermo, R0551)) was added and mixed by vortexing. Samples were incubated at -20°C overnight, then centrifuged at 21,000 g for 30 minutes at 4°C, and supernatants were discarded. Pellets were washed with 900 µl ice-cold 80% ethanol (Fisher Scientific, BP2818-4), centrifuged at 20,000 g at 4°C for 10 minutes, and supernatants were discarded. Pellets were air-dried and dissolved in 20 µl nuclease-free H_2_O.

#### E) DNase treatment of RNAs

The BoMoC polymerase used in OTTR-CLASH can utilize either DNA or RNA templates (43). Therefore, eliminating any DNA from purified RNA samples is crucial. To this end, a 25-µl DNase treatment reaction was assembled (1x RQ1 RNase-free DNase buffer (Promega), 1 U/µl murine RNase inhibitor, 0.05 U/µl RQ1 DNase (Promega # 9PIM610) and incubated at 37°C for 30 minutes.

After DNase treatment, RNA was extracted using PCI to remove the DNase. 25-µl DNase reactions were first adjusted to 300 µl with nuclease-free H_2_O, then 400 µl PCI was added to the mixture. Samples were incubated at room temperature for 8 minutes with rigorous shaking at 1400 rpm. Organic and aqueous phases were separated by centrifugation at 4°C for 10 minutes at 18,000 g. The aqueous supernatants containing the RNAs were transferred to new 1.5 nuclease-free tubes on ice.

To remove residual phenol, a further extraction with chloroform-isoamyl alcohol (IAA) (49:1; Sigma, 123-51-3 and MP Biomedicals, 155078) was performed. 300 µl of the chloroform-IAA mix was added to each RNA sample. Samples were mixed and centrifuged for 10 minutes at 18,000 g at 4°C. Upper phases were collected into new tubes on ice, and precipitated overnight after adding and mixing 700 µl of a 1:1 isopropanol-ethanol solution, 25 µl 3 M sodium acetate (pH = 5.2), and 2 µl glycogen (20 mg/ml). RNAs were collected by centrifugation at 21,000 g for 30 minutes at 4°C, and their supernatants were discarded. RNA pellets were then washed using 80% ice-cold ethanol (1 ml; (Fisher Scientific, BP2818-4)) and centrifuged at 20,000 g for 15 minutes at 4°C. Supernatants were discarded, and pellets were air-dried and then dissolved in 7 µl nuclease-free H_2_O. To help solubilize the RNAs, samples were vortexed and then incubated on ice for 15 minutes prior to storage at -80°C.

### cDNA synthesis using BoMoC polymerase

We performed OTTR-CLASH reactions using primers compatible with Illumina sequencing. The primer duplex (PD), adapter template (AT) and PCR primers are described in Supplemental Data S1. Oligonucleotide stocks for OTTR were prepared at a concentration of 100 µM concentration in water. To generate the PD duplex, the DNA and RNA strands were mixed in an equimolar ratio (18 µl of 100 µM DNA + 18 µl of 100µM RNA; 50 µM final concentration of the duplex). Strands were annealed by heating them to 72 °C for 5 min and slowly cooling down to 4 °C at a rate of - 0.2 °C/second. Thereafter, 36 µl of 100 µM AT oligonucleotide was added to the 36-µl annealed PD duplex. Finally, 928 µl nuclease-free H_2_O was added, and mixed by gentle pipetting on ice. Aliquots of 50 µl of this solution (hereafter called ‘buffer 4B’, 1.8 µM primer duplex + 3.6 µM adapter template) were frozen and kept at -80 °C until use. We avoided repeated freeze-thawing of buffer 4B and minimized its exposure to light because of the fluorophores.

The input for this OTTR reaction must be highly pure RNA (free of phenol, chloroform and other chemicals) with terminal 3’ G nucleotides (Fig. 1A). All the steps for preparation of the reaction tubes for OTTR are performed on ice. RNAs were first added to separate 0.2 ml nuclease-free tubes on ice, and water was added to a final volume of 7 µl. Then 10 µl of 2× cDNA synthesis buffer (30 mM Tris-HCl (pH = 7.0), 232 mM KCl, 8 mM MgCl_2_, 7% PEG8000, 32 mM (NH_4_)_2_SO_4_, 9.20% glycerol, 2.9 mM DTT) was added to each sample. The OTTR 2× cDNA synthesis buffer is made and stored without DTT. Just before starting each OTTR experiment, per each 190 µl aliquot of this buffer, 10 µl freshly made 0.56 mM DTT (Thermo, R0861) is added and mixed, then kept on ice.

The deoxyribonucleotides in OTTR reactions (Thermo, 10297018) are provided in a solution termed ‘buffer 4A’ (44) (4 mM dGTP, 0.8 mM dCTP, 0.8 mM dTTP, 0.04 mM dATP, 3 mM 2’-amino-dATP (dDAP-TP; APExBIO, B8079), 10 mM MgCl_2_, 900 mM KCl, 40% PEG6000; 50-µl aliquots are stored at -80°C, a working aliquot can be kept at -20°C to be used within a few months). One µl of buffer 4A was added to each OTTR reaction tube. Thereafter, each sample was mixed by pipetting up and down ten times. Then, 1 µl buffer 4B was added to each tube, without mixing or pipetting. Finally, 1 µl wild-type BoMoC enzyme (10 µM) (44) was added to each sample and the entire mix of each reaction was spun to collect all droplets from the side of the tubes to the bottoms. Reactions were then pipetted gently twenty times without making air bubbles. The 20 µl cDNA synthesis reactions were placed in a thermocycler block for 30 minutes at 37°C.

After cDNA synthesis, the RNAs were degraded. Duplexes were first denatured at 95°C for 5 minutes, then a combination of thermostable RNase H (20 mg/ml, 1 µl, (Thermo, M0523S)) and RNase A (20 mg/ml, 1 µl, NEB, T3018L) were added. After mixing, the tubes were placed back into a thermocycler and incubated for 15 minutes at 55°C. Next, proteins were degraded by adding 28 µl proteinase K solution (1 µl proteinase K (NEB, P8107S) + 27 µl cDNA stop solution: 50 mM Tris-HCl (pH 7.0), 20 mM EDTA, 0.1% v/v SDS) to each tube and incubation at 55 °C for 15 minutes in thermocycler block, followed by heating at 95 °C for 5 minutes. cDNAs were purified using Zymo Oligo Clean and Concentrate kits (Zymo D4060), eluting with 6 µl nuclease-free H_2_O into clean nuclease-free 1.5 ml tubes. An equal volume (6 µl) of 2x xylene cyanole loading dye (95% formamide, 5 mM EDTA (pH = 8.0), 0.05% w/v xylene cyanole) was added to each cDNA sample and mixed, then samples were kept at -20°C until the subsequent analysis by denaturing PAGE.

### cDNA size-selection via denaturing PAGE

OTTR-CLASH reactions generate multiple species of products (e.g. Fig. 1C). Therefore, cDNAs of interest were size-selected, via cutting and extraction from mini 8% denaturing PAGE gels (59) containing 7% urea (Sigma, 57-13-6). These gels separated reaction products with inserts from oligonucleotide dimers and smaller products (Fig. 1C and S1). Electrophoresis was monitored using xylene cyanol and bromophenol blue dyes (migrating at 76 nt and 19 nt on 8% denaturing PAGE, respectively (59)) in the 20 bp size ladder (Thermo, SM1323). The gel was run until xylene cyanol (corresponding to the size of insert-less oligonucleotide dimer products (Fig. 1C)) was two-thirds of the way to the bottom of the gel. Because of the small amounts of chimeric molecules formed, and the multiple samples analyzed on the same gel, we avoided staining the gels in a shared bath. Instead, the OTTR cDNAs were detected via the IRDye-800 on their 5’ ends. For making the molecular size marker visible, we mixed the ladder with SYBR-Gold (Invitrogen, S11494) prior to electrophoresis.

After running the gel, the gel was placed onto Saran Wrap and analyzed on a LI-COR Odyssey M imaging system, visualizing both the IRDye-800 (excitation λ = 800 nm) and the SYBR-Gold-stained ladder (excitation λ = 488 nm). Gel images were opened in ImageJ (60) and printed at actual size, with signal adjusted to display cDNA products. The gel on Saran Wrap was placed on top of and aligned with the printed image and the cDNA regions of interest were excised (Fig. 1C). The gel slices were transferred to pre-labeled tubes and crushed with a clean 1 ml pipette tip, then 500 µl cDNA elution buffer (300 mM NaCl, 10 mM Tris-HCl (pH = 8.0), 1 mM EDTA) was added and mixed by vortex. Samples were incubated at 70 °C for 1 hour to elute cDNAs into solution. Samples were centrifuged for 10 minutes at 21,000 g at 4°C and their supernatants (∼ 450 µl) were collected and transferred to new pre-chilled tubes on ice. When necessary, supernatants were again centrifuged and transferred to new tubes to remove residual gel fragments. To each eluted cDNA sample, 1 µl of 20 mg/ml glycogen (RNA grade, Thermo, R0551) was added. Samples were vortexed, split into multiple tubes, and 3 volumes of 100% ethanol were added. Any remaining gel fragments became opaque and were removed with a pipette tip. Samples were vortexed and incubated at -20 °C for 30 minutes, and the cDNAs then were pelleted by centrifugation at 21,000 g for 15 minutes at 4°C. After centrifugation, 90% of the solution was aspirated and discarded and 1 ml -20 °C-chilled 70% ethanol was added to the pellets followed by a brief vortex. The pellets were incubated at -20 °C for 15 minutes and were then centrifuged again at 21,000 g for 15 minutes. After centrifugation, nearly all the ethanol solution was aspirated and discarded. The tubes were spun briefly and all remaining liquid was carefully removed using fine micropipet tips. The remaining cDNA pellets were air-dried and resuspended in 30 µl nuclease-free water.

### Quantitative real time PCR (qPCR)

#### A) qPCR

The number of PCR cycles required to amplify libraries for high-throughput sequencing was determined experimentally via quantitative real time PCR (hereafter, qPCR)(61). 1 µl of the size-selected, purified cDNAs was analyzed in 10 µl reactions containing 0.5 µM qPCR primers (Supplemental Data S1) and 5 µl PowerUp SYBR-Green Master Mix (2×) (Applied Biosystems, A25741). Two or three technical replicates were performed, using the following amplification program: 1) Pre-incubation at 50°C for 2 minutes, 2) initial denaturation at 95 °C for 5 minutes, 3) 40 cycles amplification at 95 °C for 15 seconds and 60°C for 1 minute. Since 1 µl of purified cDNA was used in each qPCR reaction, an end-point PCR reaction on the remaining cDNA preparation (29 µl) can yield 29 times more product. Therefore, subtraction of the square-root of 29 ≈ 5.4 from the observed Ct values provides an approximate optimal cycle number for the end-point PCR.

#### B) End-point PCR for library amplification and sample indexing

Using NGS-compatible PCR primers which contain both sample barcodes (indexes) and the P5 and P7 sequences for Illumina sequencers (NEB primer sets E7335S or E7500S, Supplemental Data S1), full length libraries were amplified using the determined PCR cycle numbers. 50 μl reactions contained 25 μl Phusion High-Fidelity PCR Master Mix (2×) (NEB, M0531S), 0.5 μM of each primer, and 5 μl of the cDNA template, using the following program: Initial denaturation: 98°C for 30 seconds (1 cycle); variable cycle numbers (≤ 18 cycles) of denaturation, annealing and extension at 98°C for 10 seconds, 60°C for 30 seconds and 72°C for 30 seconds, respectively; final extension at 72°C for 10 minutes; and finally holding at 4°C.

To remove free primers and primer dimers, PCR products were purified twice using AMPure XP beads (Beckman Coulter, A63881; 1.8× reaction volume for purifying fragments ≥ 100 bp, document number: B37419AB). The purified libraries were eluted in a final volume of 25 µl nuclease-free H_2_O. After checking the size and purity of the PCR products via native 10% PAGE (Fig. 1D), concentrations of each library were determined by Qubit ((Invitrogen, Q32851) and size distributions were analyzed via Tape-Station (Agilent, G2992AA) (Supplemental Fig. S2B). Libraries were analyzed on Illumina machines with paired-end, 2 x 150 base sequencing by Azenta/Genewiz.

### Bioinformatics analysis of OTTR-CLASH data using fastChiRA

For each sample, sequencing quality of raw reads was checked using FastqQC (v0.11.9, https://github.com/s-andrews/FastQC). A quality control report for all samples of an experiment was synthesized by integrating per-sample FastQC results with MultiQC (v1.14) (62) . Adapter sequences and sequencing artifacts, polyG, of length greater than or equal 10 nt were trimmed away from R1 (3’-adapter AGATCGGAAGAGCACACGTCTGAACTCCAGTCAC) and R2 (3’-adapter AGATCGGAAGAGCGTCGTGTAGGGAAAGAGTGT) separately using Cutadapt (v4.6) (63) with the following explicit option settings: --cores 8 -a file:adapter.fa --nextseq-trim 5 --minimum-length 22 -e 0.15 --report full --overlap 10 --times 3, where adapter.fa is a FASTA file containing all 15-mers derived from the full-length adapter sequences. This trimming strategy was proposed by Manakov et al. in light of the potential existence of multiple adapter fragments (36) in chimeric eCLIP assays. The phenomenon was also confirmed in our OTTR-CLASH data. The trimmed R1 and R2 were paired up using repair.sh in the BBMap package (v39.26-0) (64), resulting matching fastq files for R1 and R2, and a mixture of single end reads of orphan R1 and R2. The re-paired R1 and R2 were merged if overlapping was detected between paired-end reads using FLASH (v1.2.11) (65), with the explicit option settings: --min-overlap 15 --max-mismatch 0.15 --max-overlap 151 -z -t 8. The merged reads were combined with orphan R1 outputted by FLASH and repair.sh to generate a new fastq file. Adapter trimming was performed on the combined reads to eliminate residual adapter sequences at the 5’ and 3’ ends using Cutadapt with the same settings. polyX at 3’ ends were removed by fastp (v1.0.1)(66) with the explicit option settings: --disable_adapter_trimming --trim_poly_g -- poly_g_min_len 10 --trim_poly_x --poly_x_min_len 10 --length_required 22. Subsequently, the resulting reads were deduplicated based on the sequence identities, which including 6-nt UMIs at the 5’ end using chira_collapse.py in a performance-enhanced version of ChiRA (67) (fastChiRA, https://github.com/haibol2016/fastChiRA, v1.4.14). For more details about the performance enhancement, see https://github.com/haibol2016/fastChiRA/blob/master/CHANGELOG.md.

To prepare the mapping references and annotation files for fastChiRA-based analyses, mature human microRNA sequences in the FASTA format and the corresponding GFF3 file were downloaded from miRBase (v22.1, https://www.mirbase.org/download/), while the primary human reference genome FASTA file and the GTF annotation file were downloaded from GENCODE (v47, https://www.gencodegenes.org/human/release_47.html) (68). The human mature microRNAs sequences were converted to DNA sequences by converting all “U” to “T” with a custom script. All microRNA-related entries were removed from the GTF file, and a reference transcriptome was generated using gffread (v0.12.7) (69) from the filtered GTF file.

Then the GFF3 file for human mature microRNAs was concatenated with the filtered GTF file to get a complete annotation file. The collapsed reads in the FASTA format were further analyzed using fastChiRA, by sequentially running the following scripts: chira_map.py, chira_merge.py, chira_quantify.py and chira_extract.py. fastChiRA outputted well-annotated chimeras, and singletons as well as summarized interactions between microRNAs and their potential targets.

### Public chimeric RNA-seq data analysis (chimeric eCLIP, CLEAR-CLIP, and qCLASH)

For each sample, sequencing quality of raw reads was assessed using FastqQC (v0.11.9, https://github.com/s-andrews/FastQC). Per-dataset quality control reports were generated using MulitQC (v1.14) (62). Public chimeric eCLIP RNA-seq dataset (GSE198251) (36) were pre-processed as described above, given that the same adapter sequences were used. Another chimeric eCLIP RNA-seq dataset (GSE297116) which were generated using the same protocol as for GSE198251 was preprocessed slightly differently because the FASTQ files downloaded from GEO had been adapter-trimmed and deduplicated (70). A close examination revealed that some deduplicated reads contain adapter sequences at 5’ ends. So, a second round of adapter trimming was performed using Cutadapt with the same option settings as above. For the CLEAR-CLIP data (32), the 3’ adapter sequence, AGTGTCAGTCACTTCCAGCGGATCTCGTATG, was trimmed using Cutadapt as above.

Additionally, 20 nt (4-nt sample indexes and adapter sequences, NNNNAGGGAGGACGATGCGG) found in all reads was trimmed using fastp with the following settings: --disable_adapter_trimming --disable_trim_poly_g --length_required 16 -- trim_front1 20 --thread 4. Based on the original publication (32), it was assumed that a NNNNG UMI exists for all reads. However, we were unable to identify such a subsequence for most reads. Therefore, we didn’t consider the UMI sequence in read deduplication for this dataset.

For the qCLASH data (71), adapter sequences and sequencing artifacts were trimmed as described for OTTR-CLASH data, except that 3’-adapter TGGAATTCTCGGGTGCCAAGGAACTC for R1 and 3’-adapter GATCGTCGGACTGTAGAACTCTGAAC for R2 were used. Other preprocessing procedures that were applied to OTTR-CLASH data were also applied to the public data where appropriate. Preprocessed public data was further analyzed using fastChiRA as described above.

Datasets analyzed are listed here:

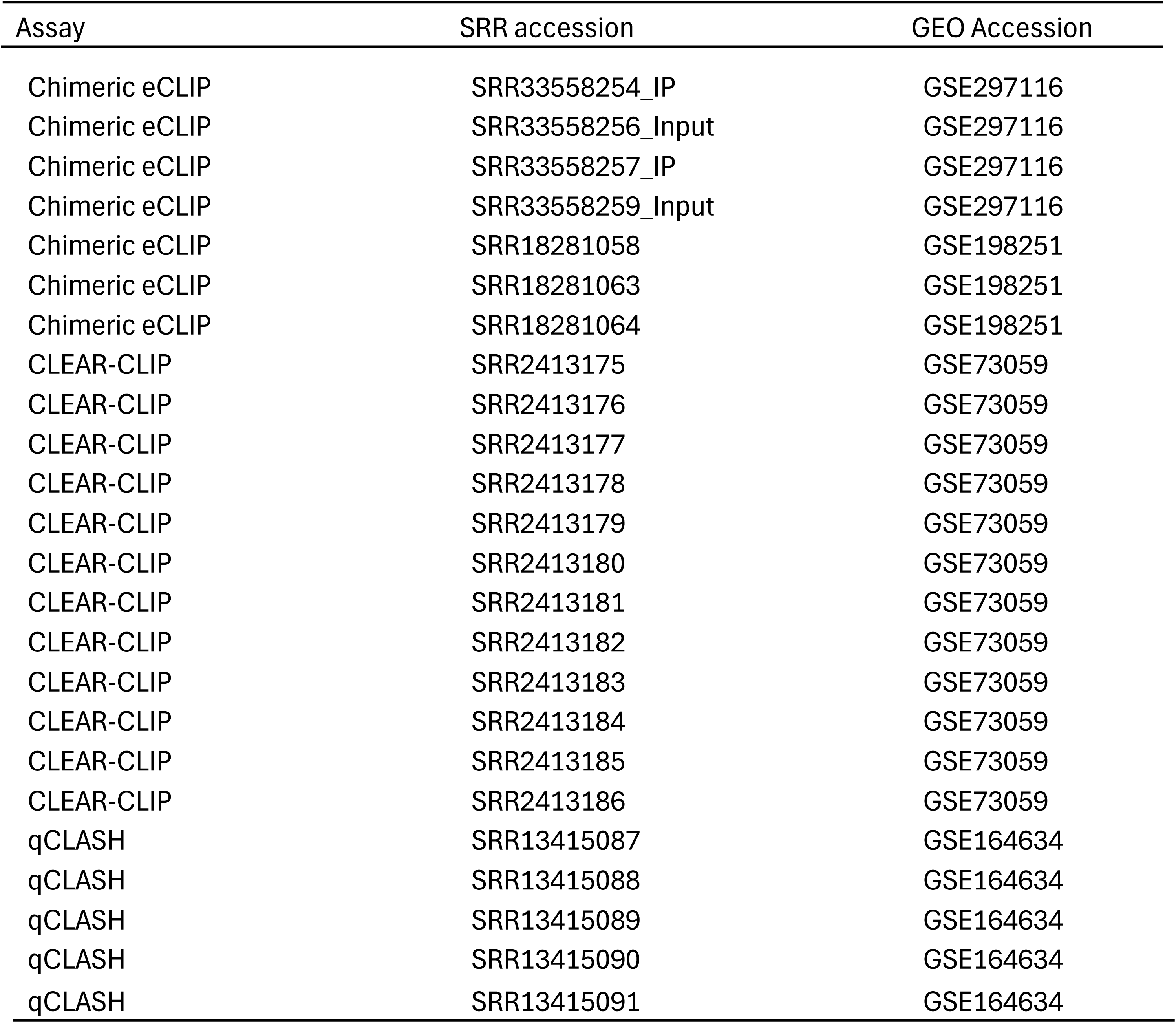

To statistically analyze the frequency of chimeras in the OTTR-CLASH and public datasets, the raw percentage data (Supplemental Data S2 and Fig. 2D) were first arcsine transformed (arcsine of the square root of the fraction of chimeric reads (72)) and analyzed via one-way ANOVA, with Dunnett’s multiple comparisons test (GraphPad Prism, v11).

**Figure 2:**
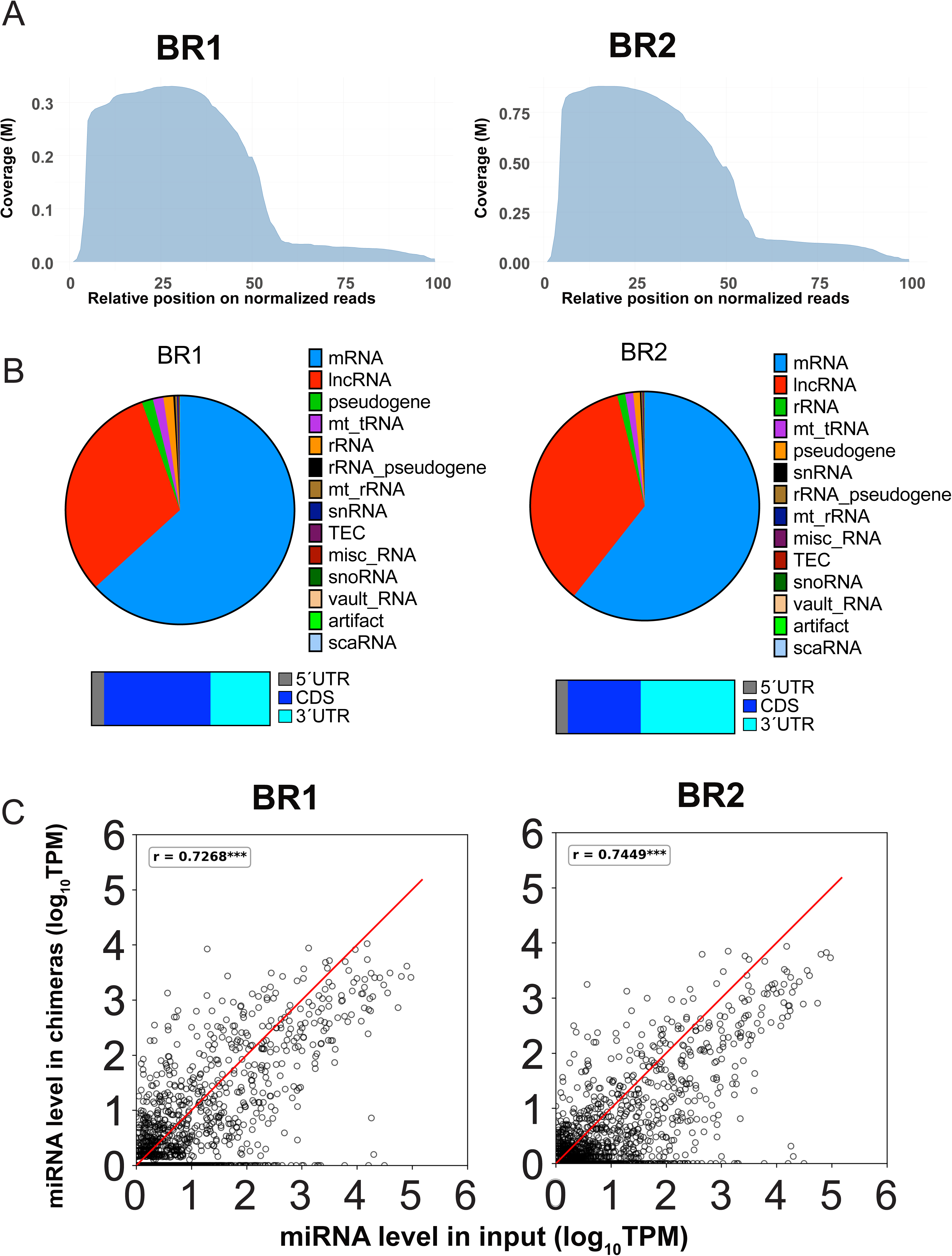
Characterization of chimeric reads from OTTR-CLASH chimeric samples BR1 and BR2. A. The location of human mature miRNAs within normalized read lengths. The enrichment of miRNAs at the 5’ end of the reads are expected for strands ligated in the context of the Ago2 complex in this protocol (Fig. 1A). B. The types of target RNAs in chimeric reads were categorized (circle plots). The most common targets are messenger RNAs (mRNAs, light blue), followed by long noncoding RNAs (lncRNAs, red). For mRNA targets, the locations within subregions (5’UTR, CDS, 3’UTR) were analyzed, with 3’UTR and CDS predominant (lower diagrams). C. Comparison of expression levels of miRNAs detected in chimeras compared to input RNA samples. These values are expected to be correlated if the OTTR-CLASH protocol is unbiased in the recovery of different sequences.

### microRNA distribution along potential chimeric reads

The U-converted human mature microRNA sequences were mapped to the collapsed reads using Bowtie (v1.3.1) (73) as described (32, 36). Alignments of microRNA to collapsed reads were filtered to only keep positive strand alignments. The start and end positions of mature microRNA on each read were determined based on the filtered alignment file and normalized against the corresponding read length for all reads of length greater than 39 nt, which were considered as candidate chimeras. The normalized start and end positions of mature microRNAs on the candidate chimeras were converted into IRanges using the R/Bioconductor package GenomicRanges (v1.58.0) (74) . Coverage was calculated from all IRanges for a sample and visualized as geom_area using R packages, ggplot2 (v3.5.2) (75) and patchwork (v1.3.1) (76).

### smallRNA-seq data analysis

Quality control of raw sequencing data was performed as above. For smallRNA-seq data generated in this study, adapter sequences were trimmed using fastp with the following explicit option settings: --adapter_sequence AGATCGGAAGAGCACACGTCTGAACTCCAGTCA -- trim_poly_g --poly_g_min_len 10 --trim_poly_x --poly_x_min_len 10 --length_required 14. For small RNA-seq data generated by Manakov et al. (36), only reads containing the adapter sequence “AACTGTAGGCACCATCAAT” followed by 12-nt UMIs were considered valid and trimmed with a custom Python script allowing one error (one substitution or indel) for matching the full adapter sequence, which also appended the UMI to the read names simultaneously.

For microRNA-only quantification, reads were mapped to U-converted human microRNA sequences using Bowtie (v1.3.1) with the following explicit settings: -v 2 -q --best -k 1 --seedlen 10 --sam --no-unal -p 8 --seed 12345, allowing up to 2 mismatches. Only reads aligned to the positive strand were kept. Reads aligned to each mature microRNA were count using a custom script, with UMI considered if available. The expression level in the TPM scale for each microRNA was calculated by normalizing per-microRNA counts against the total number of reads aligned to mature miRNAs.

To quantify all RNA species in the smallRNA-seq library, the adapter-trimmed reads were mapped to a genome sequence-decoyed human transcriptome including all cDNA sequences and noncoding RNA as well as mature microRNA sequences, and quantitated using Salmon (v1.9.0) (77), with the following explicit settings: --threads 8 --seqBias --posBias --softclip -- softclipOverhangs --biasSpeedSamp 5 -l U --validateMapping. Gene-level expression in the TPM scale were retrieved from quant.sf files generated by Salmon by tximport (v1.34.0) (78).

### Non-chimeric RNA-seq data analysis

Quality control of raw sequencing data was performed as above. Adapter sequences were trimmed using fastp with the following explicit option settings: --adapter_sequence AGATCGGAAGAGCACACGTCTGAACTCCAGTCA --adapter_sequence_r2 AGATCGGAAGAGCGTCGTGTAGGGAAAGAGTGT --trim_poly_g --poly_g_min_len 10 -- trim_poly_x --poly_x_min_len 10 --length_required 25 --thread 4. The adapter-trimmed paired-end reads were mapped to the same genome sequence-decoyed human transcriptome including all cDNA sequences and noncoding RNA as well as mature microRNA sequences as used for small RNA quantification, and quantitated using Salmon (v1.9.0) (77), with the following explicit settings: --threads 8 --seqBias --gcBias –posBias –softclip --softclipOverhangs--biasSpeedSamp 5 -l IU –validateMappings. Gene-level expression in the TPM scale were retrieved from quant.sf files generated by Salmon by tximport (v1.34.0) (78).

### Total RNA sequencing as input controls

Approximately 20 million HeLa-S3 cells were grown and pelleted as above. 0.5ml Trizol was added, and cells were homogenized via vortex for 15 sec, and incubated for 5 min at room temperature (RT). 4.5 ml ice-cold methanol was added and mixed by inverting fifteen times, followed by incubation for 10 min at RT. Samples were centrifuged at 14,000 g for 10 min at RT and the supernatant was discarded. Samples were washed with 500 µl 0.3 M guanidine hydrochloride in 95% ethanol, centrifuged at 14,000 g for 5 min and the supernatant was discarded. Samples were then washed with 200 µl freshly diluted 80% ethanol, and centrifuged at 14,000 g for 5 min at RT. Supernatants were discarded and samples were air dried for 5 min. Pellets were resuspended in 200 µl 50 mM Tris-Cl, pH 7.0, 5% SDS, 10 mM DTT, vortexed for 5 sec, and heated at 80°C for 1 hour with shaking at 500 rpm. Samples were then cooled on ice for 1-2 min, moved to RT, and 40 µl proteinase K (20 mg/ml, NEB, P8107S) was added prior to incubation at 53°C for 2 hours at 500 rpm. RNA was purified using Monarch RNA cleanup kits (10 µg) (NEB, T2030S) and eluted in 20 µl H_2_O. RNA samples were treated with 6 U DNase I (NEB, M0303S) at 37°C for 30 min and repurified using Monarch RNA cleanup kits (10 µg) (NEB, T2030S). Samples were rRNA-depleted and libraries were prepared for RNA-seq and small RNA-seq with paired-end sequencing (2×150 layout) on an Illumina NovaSeq XPlus system by Azenta/Genewiz.

### Venn Diagram analyses

Venn diagrams were drawn using the ggVennDiagram package (79).

### Data Availability

All raw sequencing data and reads processed through our pipeline are available at NIH GEO accession GSE338564.

## RESULTS AND DISCUSSION

To test whether OTTR could improve the recovery of Ago2-bound RNA chimeras, we generated whole cell extracts from suspension cultures of human HeLa-S3 cells. Lysates were treated with RNase T1 to generate target RNAs with a 3’ terminal G base, improving the resolution of the Ago2 protein footprints to be sequenced (Fig. 1A). We confirmed that we could efficiently immunoprecipitate Ago2 from these RNase T1-treated extracts (Fig. 1B), regardless of the amount of RNase T1 used. We then performed RNA ligation reactions on the Ago2 IP beads, generating a covalent link between the miRNA and target RNA strands bound by Ago2, a key step for the experimental validation of interaction pairs. RNAs purified from these ligation reactions were then subjected to OTTR reactions, and cDNA formation was analyzed via denaturing gel electrophoresis. We observed formation of slower-migrating cDNA species in IPs performed with anti-Ago2 antibodies but not control mouse IgG (Fig. 1C). Two types of larger species were detected. One migrated at the expected size of products with a single miRNA insert and were termed “Singletons”. A fainter, slower migrating band was at the expected size of products with ligated miRNA-target pairs, and these were termed “Chimeras”. In the experiment shown in Fig. 1C, we excised the Singletons and Chimeras separately from the gel. In other experiments, we excised a single gel slice containing all material larger than the oligonucleotides dimers (Supplemental Data S2A).

cDNAs were eluted from the excised gel pieces, purified and PCR amplified for sequencing. Analysis of the PCR amplification reactions confirmed that there was more material in the Ago2 IP samples than in the IgG controls (Fig. 1D and Supplemental Data S3A). In addition, the final libraries generated from the Chimera-sized material had a larger average insert size than those generated from the Singleton band (Figs. 1D, Supplemental Data S3B). Correspondingly, upon sequencing the Chimera libraries generated a subset of reads with longer lengths (Supplemental Data S3C).

We developed a customized bioinformatics workflow to analyze these data, outlined in Figure 1E. With this in hand, we analyzed the eight OTTR-CLASH datasets we had generated (Supplemental Data S2A). We observed some notable trends. First, we compared experiments performed with or without separation of cDNAs into the lower Singleton and upper Chimera bands. Libraries generated from the Chimera bands were enriched in chimeric reads, while those from the Singleton bands were highly depleted of chimeras, confirming our expectation based on gel migration. Second, libraries generated from extracts treated with different amounts of RNase T1 all produced chimeric transcripts. Because of their high levels of chimeric reads, we focused most of our attention on biological replicate Chimera samples generated from extracts treated with 1U/μl RNase T1 (Supplemental Data S2A), which we termed BR1 and BR2 for brevity. We note that these two samples displayed highly concordant levels of miRNAs and target RNAs (Supplemental Data S3D-E).

Next, we compared the frequency of recovered chimeric reads using OTTR-CLASH relative to previous methods. To make the most accurate comparisons, we used our bioinformatic pipeline to re-analyze published datasets from the three most similar previous methods that profiled miR-target interactions: CLEAR-CLIP, qCLASH and Chimeric eCLIP (Fig. 1F, Supplemental Data S2B). CLEAR-CLIP is a variant of the earlier CLASH (Crosslinking, Ligation And Sequencing of Hybrids) method, in which Ago-bound RNAs are separated by protein gel electrophoresis and recovered after transfer to nitrocellulose membranes (32). In a subsequent method, termed ‘quick CLASH’ (qCLASH), the low-yield gel and membrane transfer steps are omitted. Instead, ligated miRNA-target molecules are directly extracted from immunoprecipitated Ago complexes (34). A more recent approach, termed ‘chimeric-eCLIP’, involved revision and shortening the qCLASH workflow (36). We analyzed two different Chimeric eCLIP datasets, one generated from HCT116 cells (GSE297116) and one generated from HEK293 cells (GSE198250).

When we compared the chimeric read percentages in OTTR-CLASH experiments to those found in the public datasets (Fig. 1F), we separately considered gel-separated Chimera and Singleton samples and the unseparated “Mixed” datasets. These comparisons indicate that the Chimera samples from the OTTR-CLASH protocol produced a higher percentage of chimeric reads compared to the other sets of samples analyzed. We also examined our data for critical features expected for authentic chimeric miRNA-target pairs. First, chimeras are expected to have the miRNA on the 5’ end, i.e. be in an “miRNA-first” configuration. This is because the enzymatic steps used in library preparation favor RNA ligation between the miRNA 3’OH group and the 5’ phosphate generated on the target RNA by the phosphatase-minus PNK treatment (Fig. 1A). We observed enrichment of miRNA-first reads in the OTTR-CLASH libraries; this was true among datasets from the other methods as well, except for the Chimeric eCLIP datasets from HCT116 cells (Supplemental Data S3D). We next categorized the type of RNA species found as targets in the chimeric reads (Fig. 2B and Supplemental Data S4A). Consistent with other studies of Ago-associated targets, the majority of RNAs found were mRNAs and lncRNAs, with mRNA interactions much more frequent in the 3’ UTR and coding sequence (CDS) regions than in 5’ UTRs (36, 70). Finally, we expected that if OTTR-CLASH had successfully captured a wide variety of miRNA-target interactions with a minimum of sequence bias, we should observe a linear relationship between the frequency of detecting a miRNA in chimeric reads and its abundance in whole cell extracts. Indeed, that is what we observed (Fig. 2C and Supplemental Data S4B). Together, these characteristics suggested that OTTR-CLASH had efficiently captured relevant miRNA-target interactions.

We then explored how specific interactions detected by OTTR-CLASH compared to those detected previously. For these analyses, we focused on the miRNA-target interactions found in the two 1U/ml RNase T1 biological replicate experiments, BR1 and BR2. We performed two separate comparisons using two types of data. First, we compared reads found in OTTR-CLASH to the four groups of public datasets described above (Supplemental Data S2B). A Venn diagram depicting the overlaps among the five methods showed minimal overlap, with the largest number of reads specific to each method (Fig. 3A). The high degree of unique interactions is most likely due to the fact that each dataset was generated with a different cell line. We next compared the OTTR-CLASH output to public databases that amalgamate lists of miRNA-target interactions: miRNATIP (80), DIANA-microT (81), miRTarCLASH (82), miRTarBase (83), and TarBase (84). Here, there was much more extensive overlap of the OTTR-CLASH data with the other classes examined, but there were still more than 87,000 interactions specific to the OTTR-CLASH data (Fig. 3B). To find the best candidates for novel interactions, we then selected those interactions found in both of the comparisons, and then filtered that list of 61,085 interactions for those found in both BR1 and BR2, resulting in a list of 1,643 candidate novel interactions (Supplemental Data S5).

**Figure 3:**
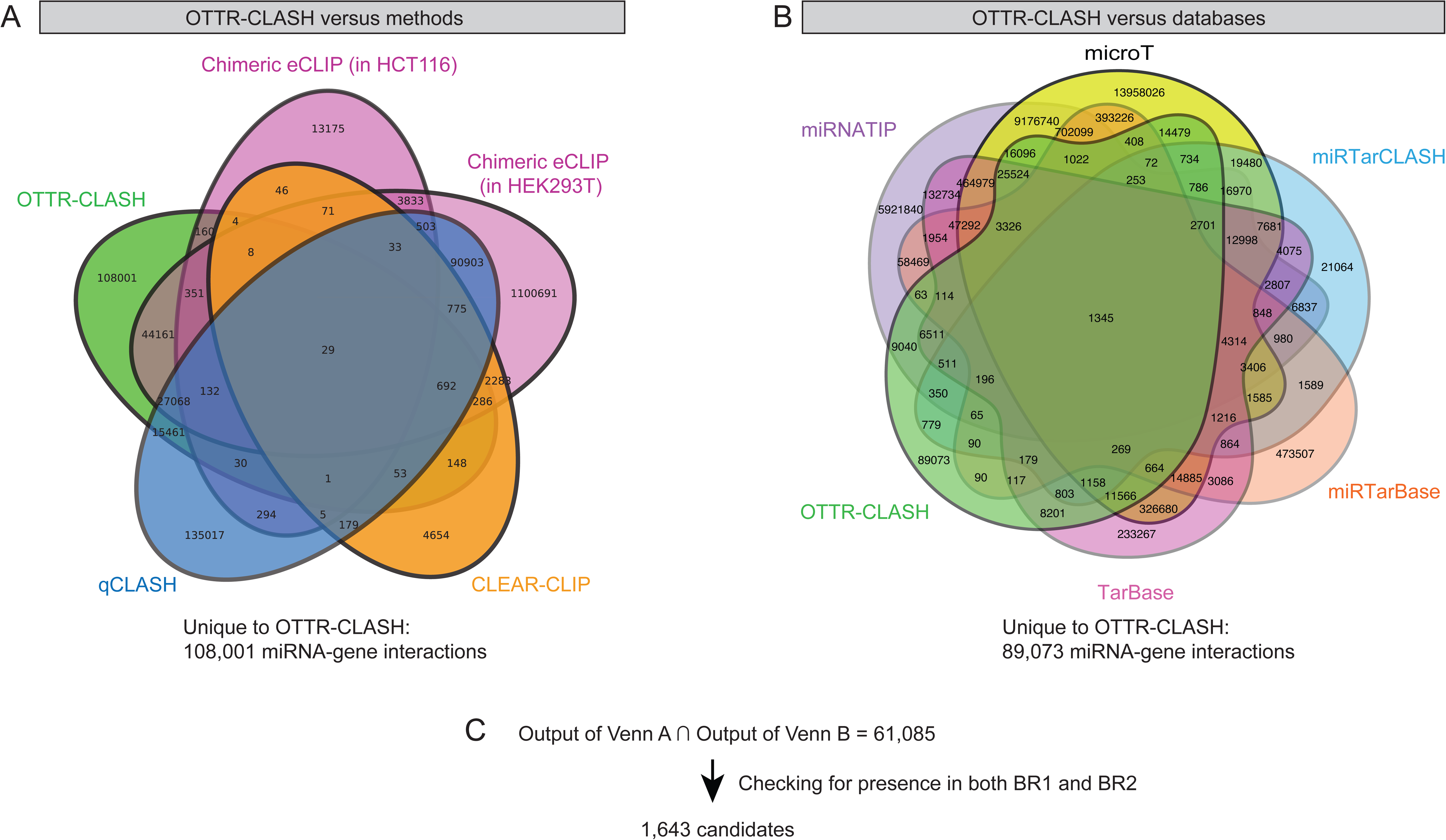
Venn Diagram comparisons of OTTR-CLASH to other methods and databases. (A) All miRNA-gene interactions detected by OTTR-CLASH (n = 10 samples) were compared to all such interactions detected by three other methods. These included one qCLASH, one CLEAR-CLIP, and two chimeric eCLIP experiments. 108,001 miRNA-gene interactions specific to OTTR were found. (B) In parallel, all miRNA-gene interactions detected by OTTR-CLASH were intersected with five miRNA-target databases (miRNATIP, microT, miRTarCLASH, miRTarBase, and TarBase). (C) The intersection between OTTR-specific interactions in panels A and B contains 61,085 novel miRNA-gene interactions. Of those, 1,643 were observed in both BR1 and BR2.

To examine these candidates in more detail, we filtered them by more stringent criteria. First, the modified ChiRA analysis in our bioinformatic pipeline calculates a minimum free energy (mfe) for the hybridization between the miRNA and target strands. We considered interactions with a hybridization energy score of < -7 kcal/mol and that were detected at least two reads per biological replicate experiment. Seventeen interaction pairs met these criteria. We disregarded one because the sequenced reads were mostly in the “miRNA-last” configuration (85). To explore the biological relevance of the remaining sixteen interactions, we consulted the Encori/Starbase database (86). This database takes miRNA and mRNA expression data from the The Cancer Genome Assembly (TCGA) (https://www.cancer.gov/tcga) to calculate co-expression frequencies across 32 types of cancer. We sought interactions for which there were at least three cancer subtypes that displayed co-variation between the miRNA and the target with p-values < 1 e^-4^, and for which the direction of regulation (that is, repressive or activating) was in the same direction in all such cancer subtypes.

Four of these novel interactions displayed these characteristics. For three of these, higher miRNA expression correlated with lower mRNA levels (Fig. 4A-C). This is the expected direction of regulation, because miRNAs frequently stimulate degradation of target RNAs via the RISC complex (13). However, we also observed one example in which higher miR-1247-5p levels correlated with greater target TM4SF1 mRNA levels (Fig. 4D). This type of positive correlation can result from direct transcriptional gene upregulation by miRNAs (87) or when miRNAs block access to the target by RNA degradation enzymes (88). We note that in this case, the miRNA target site is very near the mRNA polyadenylation site on the 3’UTR, and this mRNA undergoes regulated alternative polyadenylation (89). Therefore, the positive relationship between miR-1247-5p and TM4SF1 mRNA levels could result from upregulation of specific mRNA isoforms. We thus conclude that OTTR-CLASH can provide evidence for novel miRNA-target interactions that can be implicated in complex regulation patterns.

**Figure 4.**
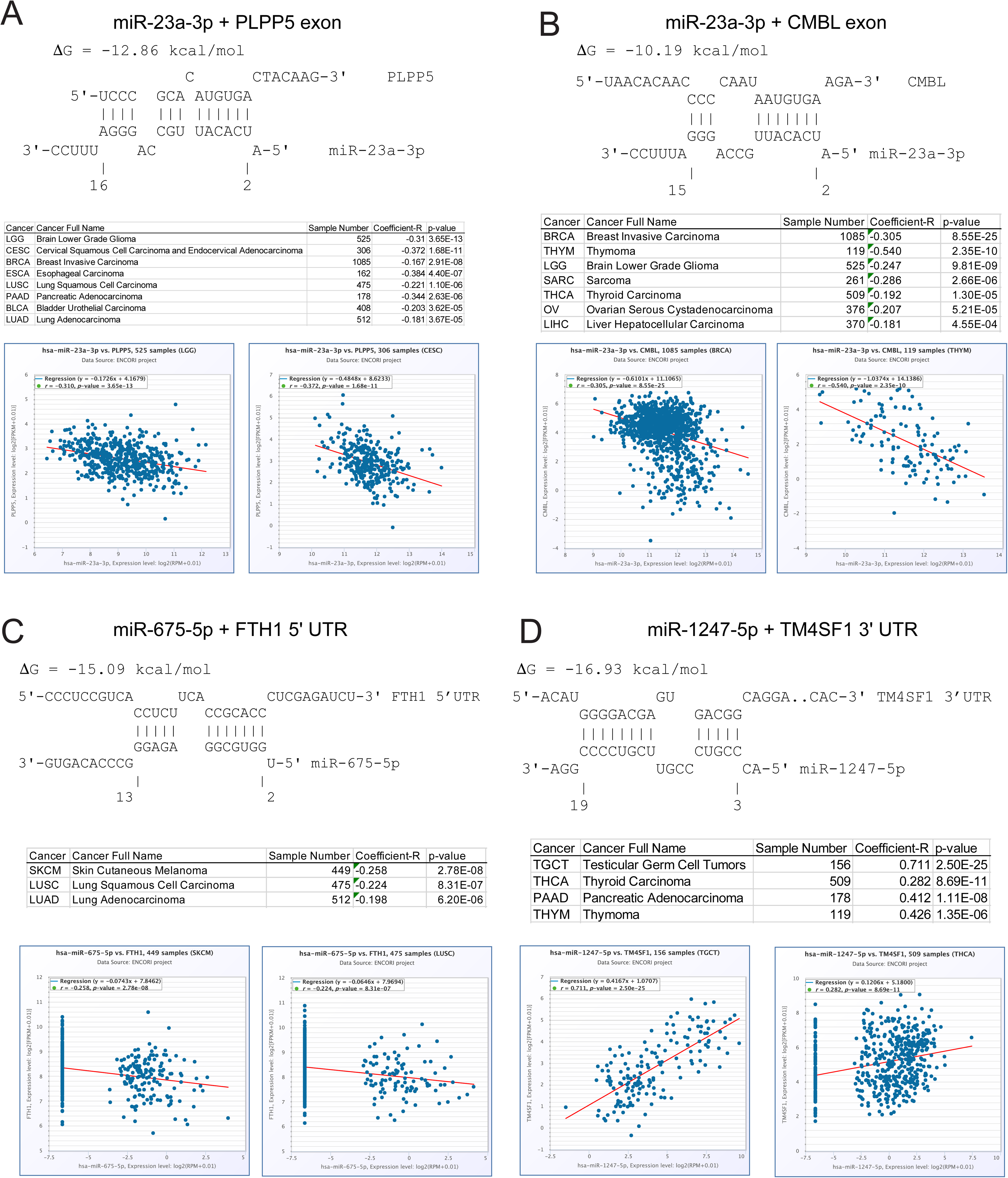
Analysis of novel miRNA-target interactions detected by OTTR-CLASH. Shown are examples of miR-target interactions detected by OTTR-CLASH that were not previously seen in public databases. For each example, the following are shown: (top) RNA hybridization and energetics predicted by IntaRNA; (middle) Data for all cancer subtypes from Starbase with p < 0.001. Listings are sorted by p-values; (bottom) Specific examples of correlations between miRNA and mRNA levels. A. miR-23a-3p/PLPP5 exon B. miR-23a-3p/CMBL exon C. miR-675-5p/FTH1 5’UTR D. miR-1247-5p/TM4SF1 3’UTR

In sum, we have presented an improved method for the discovery and analysis of Argonaute 2-associated miRNA-target RNA interactions. Our data indicates that gel fractionation of the OTTR cDNA synthesis reactions generates libraries with greater percentages of chimeric reads than non-size-fractionated samples or libraries from previous methods (Fig. 1F, Supplemental Data S2). Therefore, use OTTR-CLASH reactions that include the gel fractionation step decreases the amount of sequencing required for analysis of Ago2-associated RNAs.

UV-crosslinking is considered superior for the studies of RNA binding proteins because as a “zero-length” crosslinker, it captures direct contacts at interaction sites (90). A drawback, however, is that UV can leave bulky adducts during the deproteinization steps required for deep sequencing analyses, resulting in truncated cDNA formation (91). To avoid this problem, we utilized OTTR technology, based on a silkworm non-LTR retrotransposon reverse transcriptase that efficiently polymerizes across modified nucleotides and simultaneously joins the product cDNAs to sequencing adapters, bypassing the need for separate and inefficient ligation steps (43, 44). The improved rate of obtaining chimeric RNAs (Fig. 1D) illustrates the success of the OTTR-CLASH method. Therefore, this is another example of the superiority of this enzyme for the analysis of rare and/or chemically modified RNAs; previous examples included ribosomal footprints (45) and tRNAs (46). We envision that analogous adaptation of OTTR to other types of experiments with very low concentrations of modified input RNAs could result in similar benefits.

## Author Contributions

PDK and TP conceptualized the project and acquired funding, with investigation by PDK. Software was developed by HL, with data curation by HL and KH; all bioinformatic analyses were supervised by LJZ. Critical resources, protocols and experimental expertise were provided by LF and KC. The original draft of the paper was written by PDK and TP with review and editing from all the authors.

## Acknowledgements

We thank Hans Tobias Gustaffson and Oliver J. Rando for experimental insights, reagents and advice.

---

**Supplemental Data S1.**
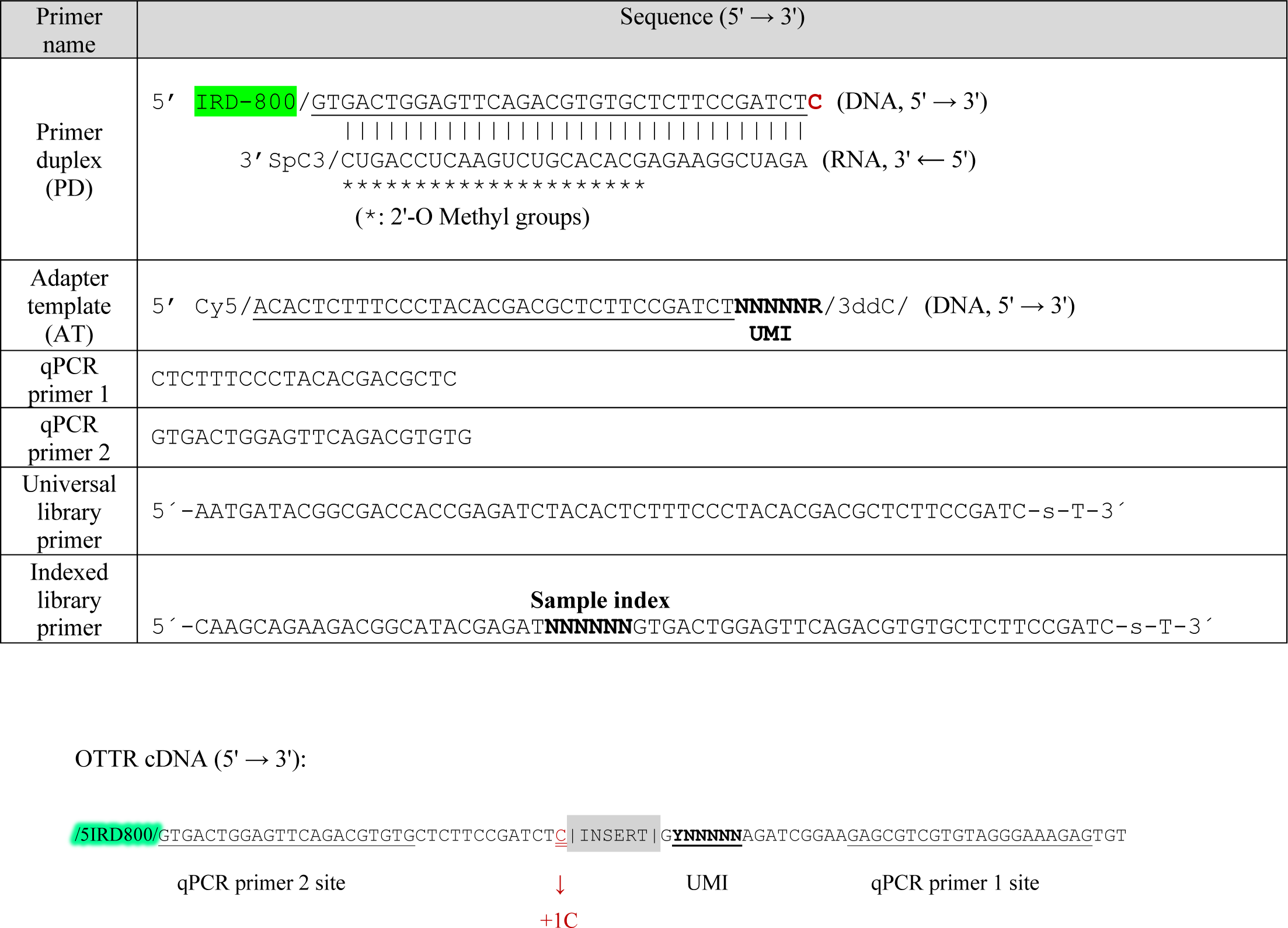
Oligonucleotides for OTTR-CLASH. The Primer Duplex (PD) is a DNA/RNA duplex with an infra-red label (IRD-800) on the 5’ end of the DNA strand and a blocked 3’ end (SpC3) on the RNA strand. The BoMoC enzyme uses the terminal 3’ C residue on the DNA strand (red, bold) to anneal to RNA templates with 3’ G bases. PD carries the sequence of the Illumina read 2 sequencing primer (underlined). The Adapter Template (AT) contains Illumina read 1 sequencing primer sequences (underlined) as well as the UMI (bold face nucleotides). It is Cy5-labeled at its 5’ end and blocked by dideoxycytidine (ddC) at its 3’ end. The structure of the cDNAs resulting from the OTTR reaction are cartooned at the bottom. The qPCR primers are used for quantitation of the OTTR cDNAs. The universal and indexed PCR primers are used to generate the final libraries for deep sequencing.

**Supplemental Data S2:**
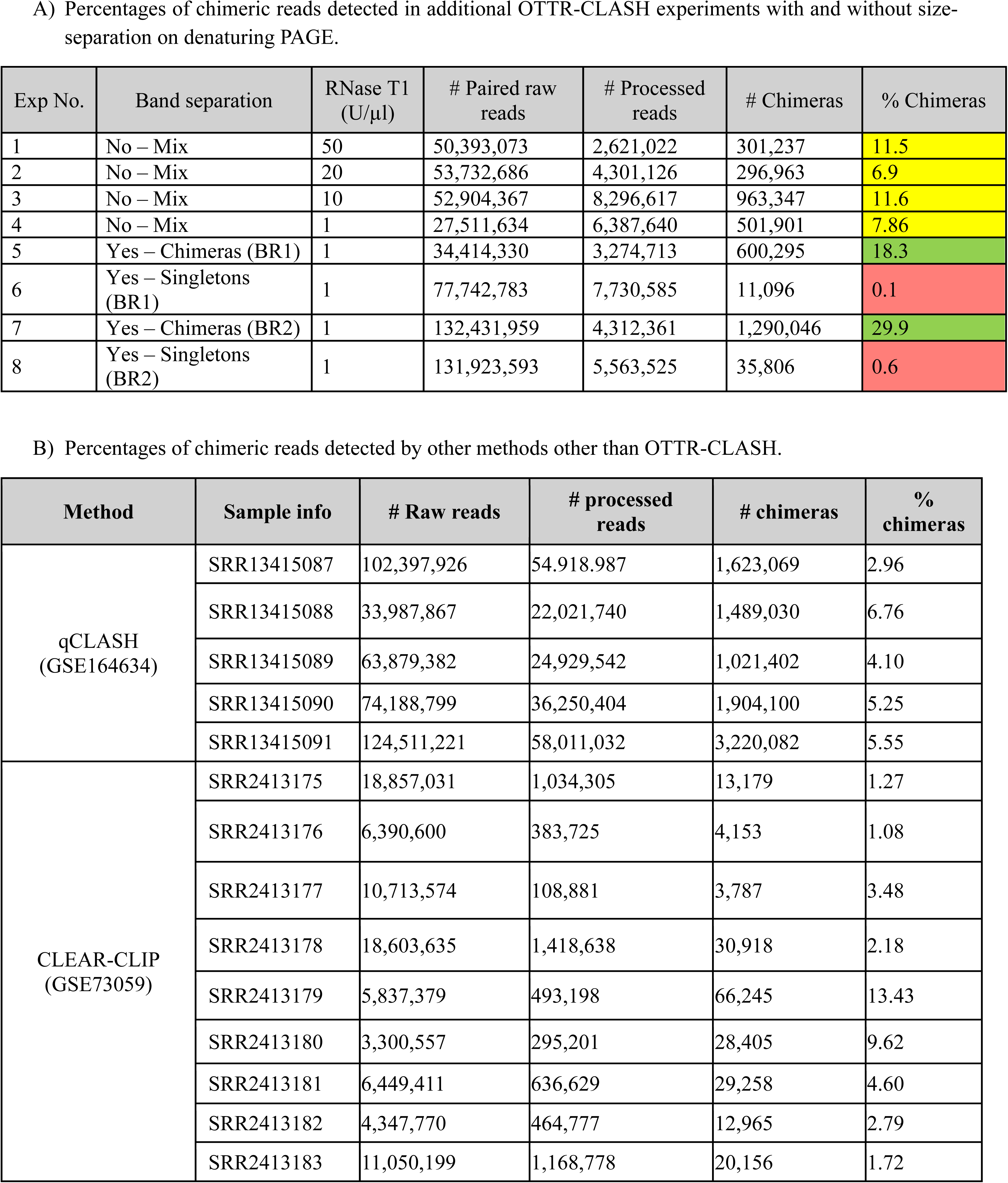
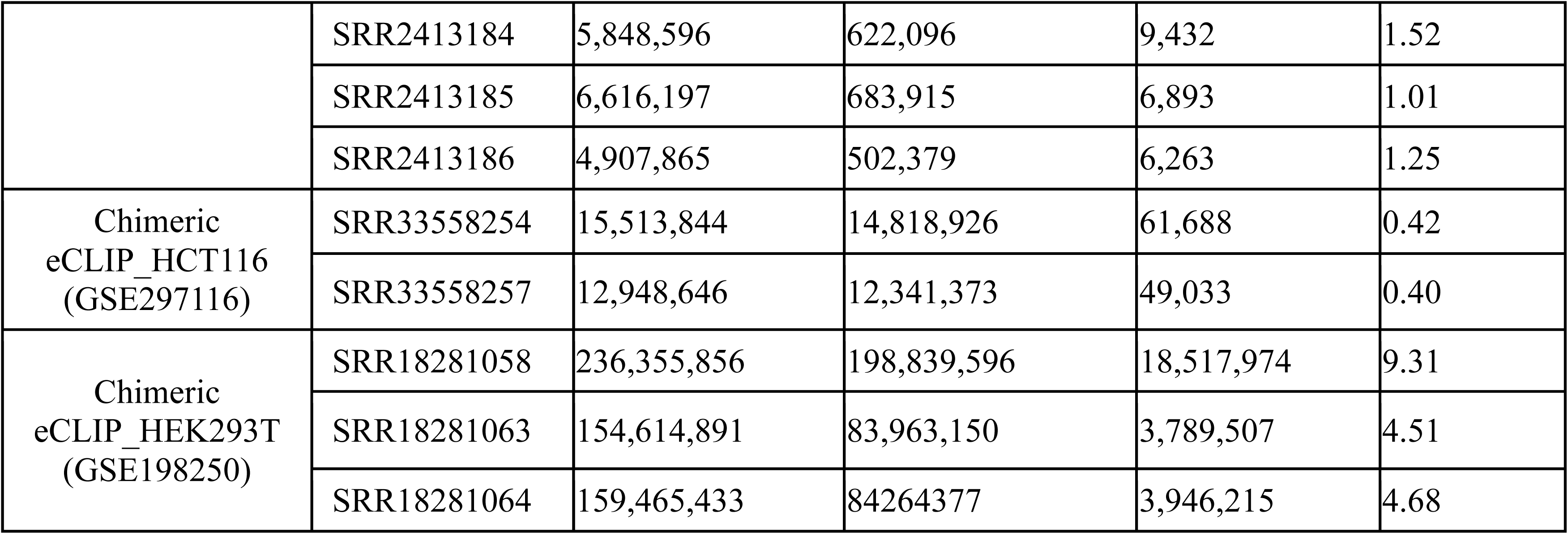
Percentages of chimeric reads in OTTR-CLASH and public datasets. A) Data from OTTR-CLASH experiments, with and without size-separation by denaturing gel electrophoresis. Data from the column on the right are graphed in Figure 1F. Yellow indicates samples that were not separated into chimeras and singletons. Green are the size-selected chimera samples, and red are the size-selected singleton samples. B) Data from public datasets obtained with other methods.

| Exp No. | Band separation | RNase T1 (U/μl) | # Paired raw reads | # Processed reads | # Chimeras | % Chimeras |
| --- | --- | --- | --- | --- | --- | --- |
| 1 | No – Mix | 50 | 50,393,073 | 2,621,022 | 301,237 | 11.5 |
| 2 | No – Mix | 20 | 53,732,686 | 4,301,126 | 296,963 | 6.9 |
| 3 | No – Mix | 10 | 52,904,367 | 8,296,617 | 963,347 | 11.6 |
| 4 | No – Mix | 1 | 27,511,634 | 6,387,640 | 501,901 | 7.86 |
| 5 | Yes – Chimeras (BR1) | 1 | 34,414,330 | 3,274,713 | 600,295 | 18.3 |
| 6 | Yes – Singletons (BR1) | 1 | 77,742,783 | 7,730,585 | 11,096 | 0.1 |
| 7 | Yes – Chimeras (BR2) | 1 | 132,431,959 | 4,312,361 | 1,290,046 | 29.9 |
| 8 | Yes – Singletons (BR2) | 1 | 131,923,593 | 5,563,525 | 35,806 | 0.6 |

B) Percentages of chimeric reads detected by other methods other than OTTR-CLASH.
| Method | Sample info | # Raw reads | # processed reads | # chimeras | % chimeras |
| --- | --- | --- | --- | --- | --- |
| qCLASH (GSE164634) | SRR13415087 | 102,397,926 | 54,918,987 | 1,623,069 | 2.96 |
|  | SRR13415088 | 33,987,867 | 22,021,740 | 1,489,030 | 6.76 |
|  | SRR13415089 | 63,879,382 | 24,929,542 | 1,021,402 | 4.10 |
|  | SRR13415090 | 74,188,799 | 36,250,404 | 1,904,100 | 5.25 |
|  | SRR13415091 | 124,511,221 | 58,011,032 | 3,220,082 | 5.55 |
| CLEAR-CLIP (GSE73059) | SRR2413175 | 18,857,031 | 1,034,305 | 13,179 | 1.27 |
|  | SRR2413176 | 6,390,600 | 383,725 | 4,153 | 1.08 |
|  | SRR2413177 | 10,713,574 | 108,881 | 3,787 | 3.48 |
|  | SRR2413178 | 18,603,635 | 1,418,638 | 30,918 | 2.18 |
|  | SRR2413179 | 5,837,379 | 493,198 | 66,245 | 13.43 |
|  | SRR2413180 | 3,300,557 | 295,201 | 28,405 | 9.62 |
|  | SRR2413181 | 6,449,411 | 636,629 | 29,258 | 4.60 |
|  | SRR2413182 | 4,347,770 | 464,777 | 12,965 | 2.79 |
|  | SRR2413183 | 11,050,199 | 1,168,778 | 20,156 | 1.72 |

**Supplemental Data S2**
|  |  |  |  |  |  |
| --- | --- | --- | --- | --- | --- |
|  | SRR2413184 | 5,848,596 | 622,096 | 9,432 | 1.52 |
|  | SRR2413185 | 6,616,197 | 683,915 | 6,893 | 1.01 |
|  | SRR2413186 | 4,907,865 | 502,379 | 6,263 | 1.25 |
| Chimeric<br>eCLIP_HCT116<br>(GSE297116) | SRR33558254 | 15,513,844 | 14,818,926 | 61,688 | 0.42 |
|  | SRR33558257 | 12,948,646 | 12,341,373 | 49,033 | 0.40 |
| Chimeric<br>eCLIP_HEK293T<br>(GSE198250) | SRR18281058 | 236,355,856 | 198,839,596 | 18,517,974 | 9.31 |
|  | SRR18281063 | 154,614,891 | 83,963,150 | 3,789,507 | 4.51 |
|  | SRR18281064 | 159,465,433 | 84264377 | 3,946,215 | 4.68 |

**Supplemental Data S3:**
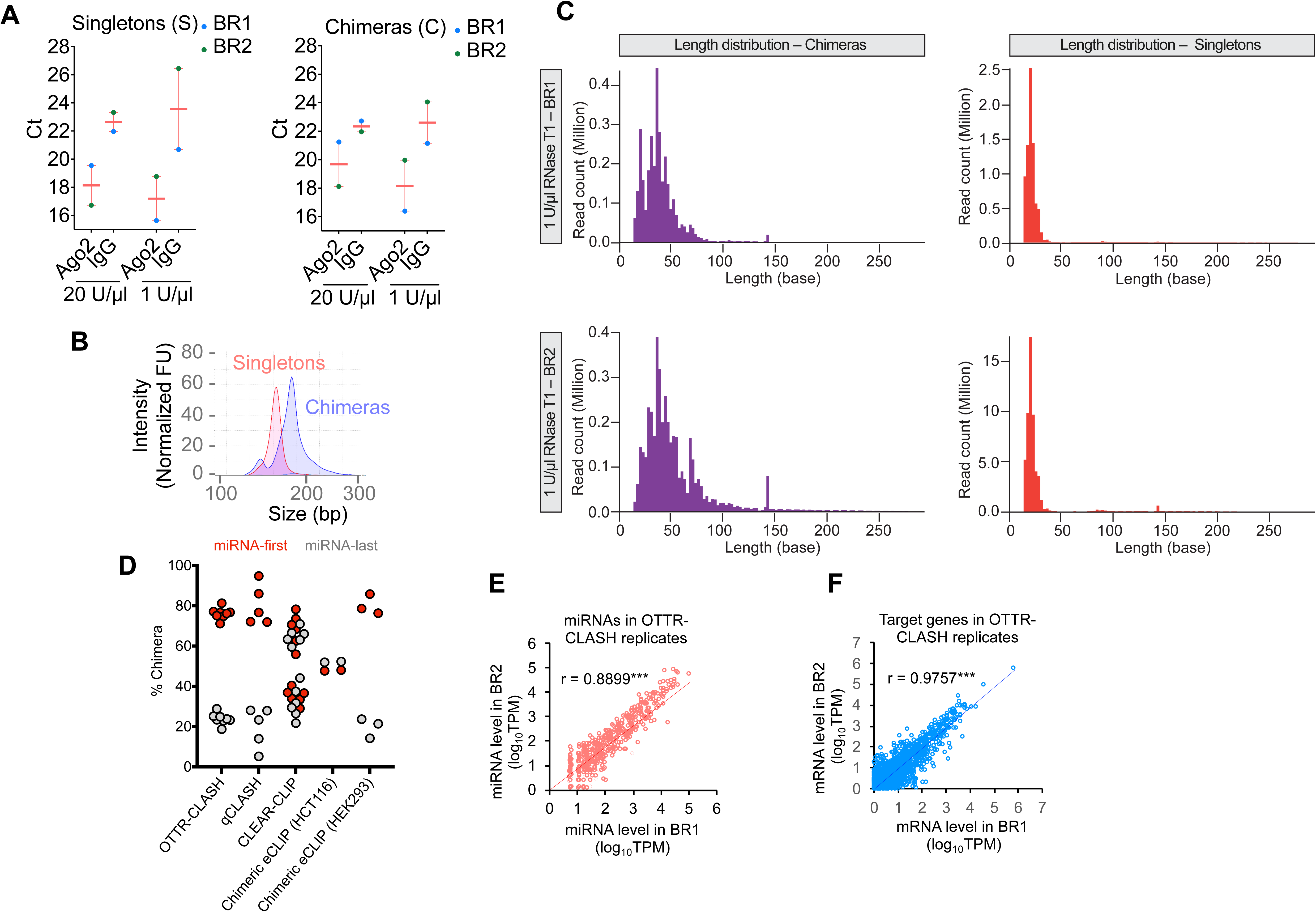
Characterization of additional OTTR-CLASH reactions. A) qPCR Ct values for the libraries analyzed in Figure 1D. The smaller Ct values in the Ago2 IP samples compared to the IgG controls indicates capture of Ago2-bound RNAs. B) An example of Tape-Station analysis of PCR-amplified libraries for precise assessment of their size distributions. Chimera products displayed a larger size distribution than singleton products. C) Sequencing read length distributions of the indicated samples. D) Percentage of chimeric reads with miR-first (red dots) or miR-last orientations (grey dots) from the indicated datasets. E) Comparison of levels of miRNA levels in the replicate OTTR-CLASH datasets BR1 and BR2. F) Comparison of levels of target mRNA levels in the replicate OTTR-CLASH datasets BR1 and BR2.

**Supplemental Data S4.**
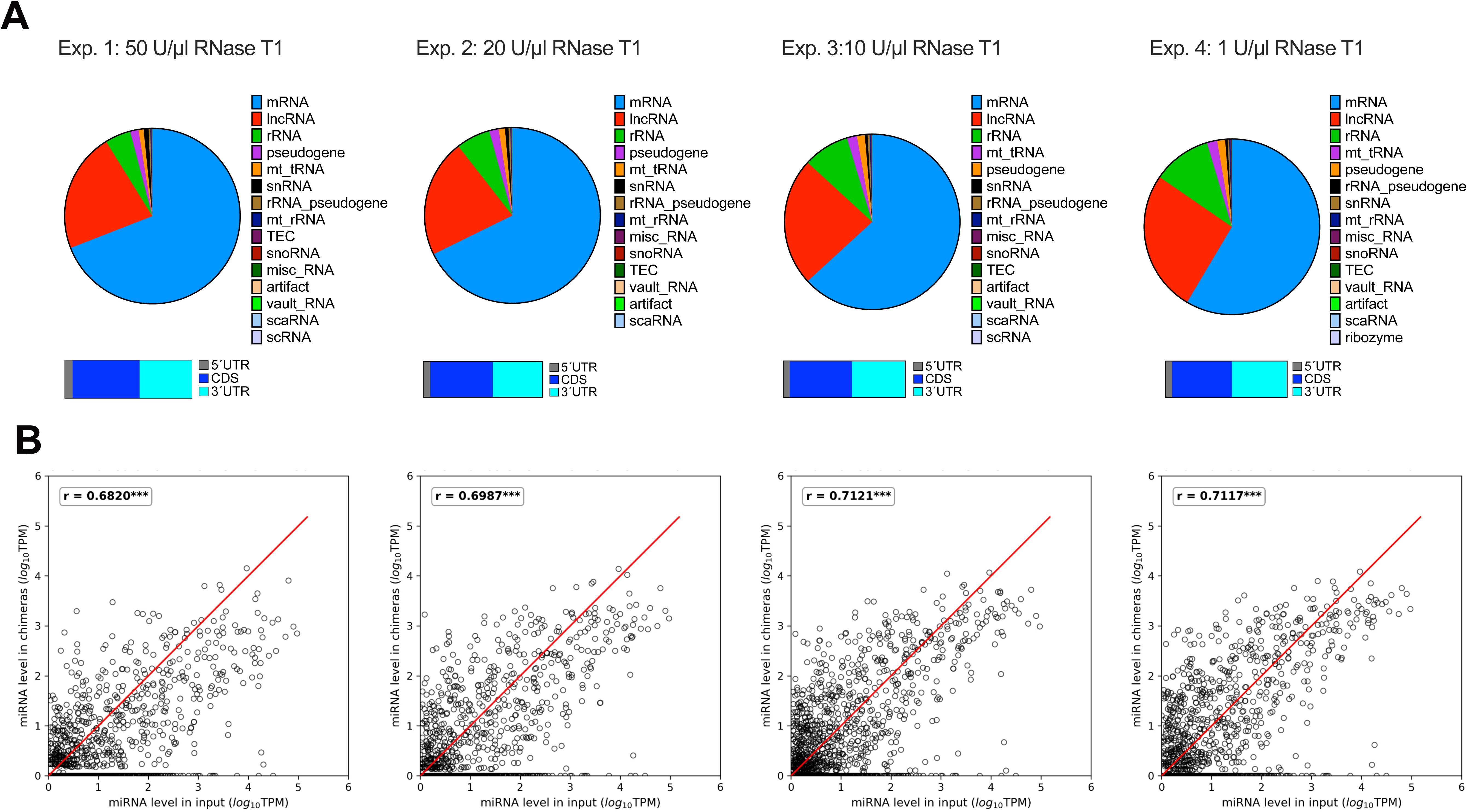
Data characterization for additional OTTR-CLASH samples. Experiment numbers (listed in Supplemental Data S1) are indicated at the top. A. Target types for additional OTTR-CLASH reactions, as in Figure 2B. B. Input/chimera recovery correlations for additional OTTR-CLASH samples, as in Figure 2C.

Supplemental Data S5: OTTR-unique candidates corresponding to the final output file of Figure 3C.

